# Cocaine Seeking And Taking Are Oppositely Regulated By Dopamine

**DOI:** 10.1101/2023.04.09.536189

**Authors:** Lauren M. Burgeno, Ryan D. Farero, Nicole L. Murray, Marios C. Panayi, Jennifer S. Steger, Marta E. Soden, Scott B. Evans, Stefan G. Sandberg, Ingo Willuhn, Larry S. Zweifel, Paul E. M. Phillips

## Abstract

In some individuals, drug-associated cues subsume potent control of behavior, such as the elicitation of drug craving^1–3^ and automatized drug use^4^. The intensity of this cue reactivity is highly predictive of relapse and other clinical outcomes in substance use disorders^5, 6^. It has been postulated that this cue reactivity is driven by augmentation of dopamine release over the course of chronic drug use^7^. Here we carried out longitudinal recording and manipulation of cue-evoked dopamine signaling across phases of substance-use related behavior in rats. We observed a subset of individuals that exhibited increased cue reactivity and escalated drug consumption, two cardinal features of substance use disorders. In these individuals, cue-evoked phasic dopamine release underwent diametrically opposed changes in amplitude, determined by the context in which the cue is presented. Dopamine evoked by non-contingent cue presentation increased over drug use, producing greater cue reactivity; whereas dopamine evoked by contingent cue presentation decreased over drug use, producing escalation of drug consumption. Therefore, despite being in opposite directions, these dopamine trajectories each promote core symptoms of substance use disorders.

## Introduction

Drug addiction is a neuropsychiatric disorder characterized by cycles of irrepressible drug use, compulsive drug seeking, and high propensity to relapse following periods of abstinence^8^. Key to these states are drug-associated stimuli (cues). When drug seeking actions are successful, individuals learn about stimuli that are reliably paired with drug receipt through classical (Pavlovian) conditioning^3, 9–11^. Importantly, these cues acquire the ability to modify behavior when they are encountered outside the usual pairing context–a process known as Pavlovian-to-instrumental transfer^12, 13^. For example, following chronic drug use in some individuals, a drug cue that was previously received as a result of drug-seeking actions (contingent presentation) can elicit drug craving^1–3^ and automatized drug-seeking^4^ when experienced in other situations (non-contingent presentation).

Many contemporary theories of substance use disorders propose that enduring changes in mesolimbic dopamine transmission underlie aberrant behaviors^14^. For example, the premise of the incentive sensitization theory is that dopamine release evoked by drug-associated environmental stimuli *increases* over the course of drug use to precipitate drug craving^7^. However, during active cocaine taking, rapid dopamine responses to drug-paired stimuli progressively *decrease* over the course of chronic drug use^15^. While these data superficially appear to be incompatible with the theory, it is important to recognize that they relate to different contingencies of stimulus presentation: Incentive sensitization pertains to reactivity to non-contingent stimulus presentation, rather than stimuli when they are experienced following drug-taking actions^15^. Importantly, drug-paired stimuli can serve different purposes depending upon their contingencies. When a drug-paired stimulus is presented in response to a drug-taking action, it acts as a feedback signal indicating that drug delivery is imminent and no further action is required, whereas the same stimulus, presented non-contingently outside the active drug-taking context, acts as a cue (eliciting stimulus) to promote drug seeking.

Here we examined how dopamine responses, evoked by non-contingent presentation of cues, evolve over the course of chronic drug use and withdrawal. We observed increasing dopamine release, a trajectory that is diametrically opposed to changes in dopamine evoked by the same stimulus when presented in response to a drug-taking action, measured in the same subjects. These opposing changes in dopamine mediate distinct hallmark features of addiction, with *decreased* dopamine to response-contingent stimuli producing increased drug consumption, and *increased* dopamine to non-contingent cues producing craving.

## Results

### Drug-use history impacts non-contingent CS-elicited phasic dopamine transmission

Male Wistar rats, implanted bilaterally with carbon-fiber microelectrodes in the nucleus accumbens core (NAcc) and a jugular catheter, were trained to receive intravenous cocaine in daily one-hour sessions (short-access, ShA; Figure 1a). Behavioral chambers were outfitted with two nose-poke ports, of which one was designated for drug-taking (side counterbalanced across animals); a house light and white noise signaled when the drug-taking port was active. A cocaine infusion (0.5 mg/kg) paired with a separate audiovisual conditioned stimulus (CS, nose-poke light and tone; Figure 1b) was delivered following a nose-poke response in the active drug-taking port (responses in the other port had no programmed consequences). Once animals reached the acquisition criterion (>10 responses in three consecutive sessions), they received an additional five baseline ShA sessions (Figure 1c). Immediately prior to the last of these sessions, non-contingent CS probe sessions were conducted with the animal in the drug-taking chamber but outside the usual drug-taking context (nose-poke ports were inaccessible, and house light and white noise were off). During each probe session, the CS was presented twice, independent of the animal’s behavior, and without drug delivery. A control group of animals, naïve to cocaine self-administration, also received probe sessions. Dopamine transients evoked by the non-contingent CS were measured with fast-scan cyclic voltammetry (FSCV) in both groups. As previously shown following comparable self-administration training^16^, the CS acquires the capacity to elicit phasic dopamine release in the NAcc when encountered unexpectedly (Figure 1d). These responses were significantly greater in animals with self-administration experience than in naïve animals (Unpaired t-test: t=2.313, df=21, p=0.03; Figure 1d), confirming that these are learned, rather than innate responses.

**Figure 1.**
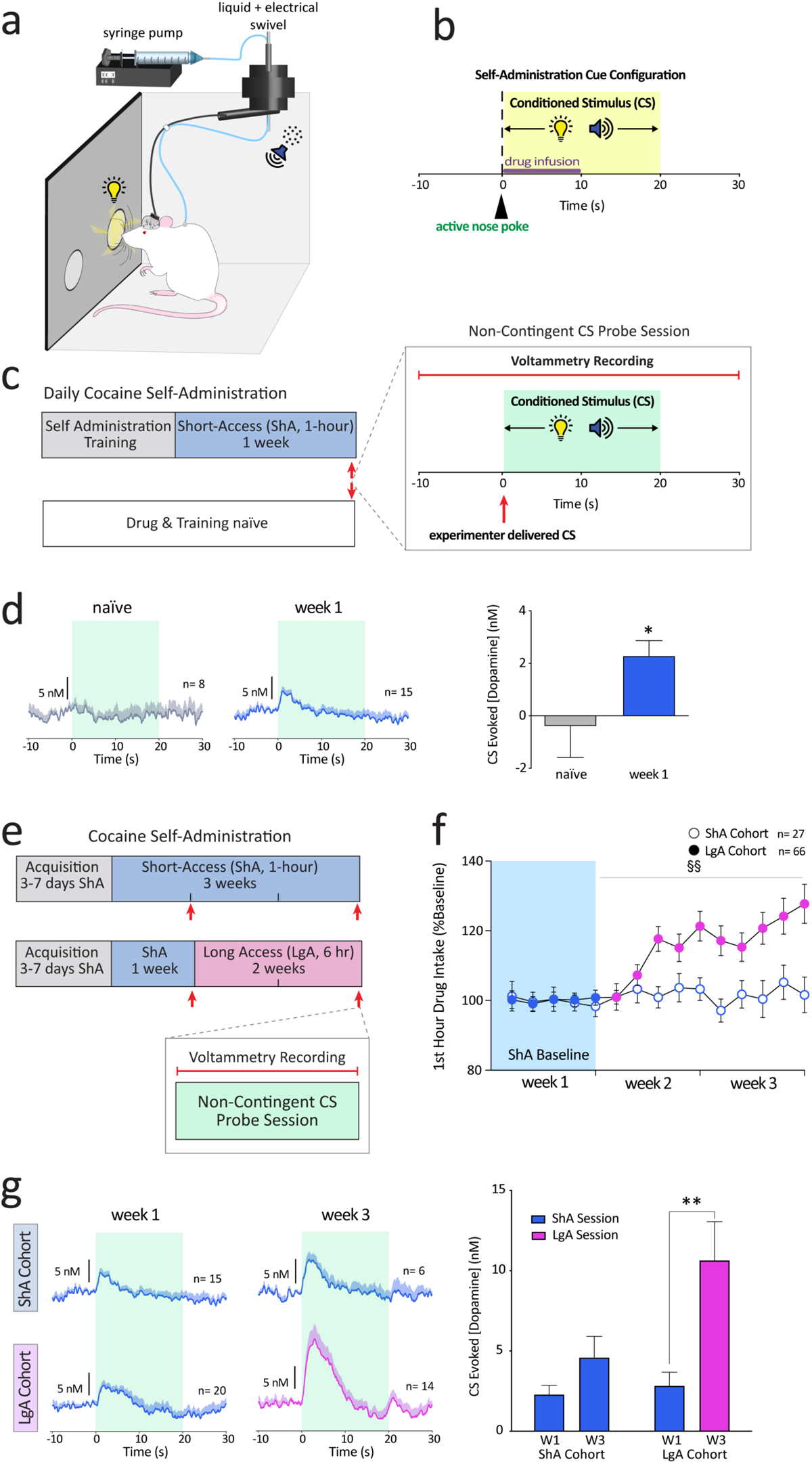
Drug-use history impacts non-contingent CS-elicited phasic dopamine transmission. **a)** Operant chamber used for cocaine self-administration. **b)** Self-administration cue configuration. Nose pokes into the active port (black triangle & dotted line) triggered intravenous delivery of cocaine (0.5mg/kg) and concurrent presentation of a 20 s audiovisual conditioned stimulus (CS; yellow box indicates CS duration). **c)** Experimental paradigm used to measure non-contingent CS evoked dopamine release. Following acquisition of self-administration behavior, subjects received an additional week of short-access (ShA; 1-hour daily) and non-contingent probe sessions (red arrows) were administered to both trained and drug naive groups. Probe sessions consisted of two CS presentations, independent of the animal’s behavior and without drug delivery, spaced three minutes apart. Cyan box indicates non-contingent CS duration. CS audiovisual properties were identical to those in a. Dopamine responses to non-contingent CS were measured with Fast-Scan Cyclic Voltammetry (FSCV). **d)** Mean (+SEM) non-contingent CS-evoked dopamine response traces from naïve controls (gray trace, left) and week 1 ShA (blue trace, middle). Right: Mean (+SEM) CS-evoked [Dopamine](nM). CS-evoked dopamine responses were larger in animals with self-administration experience than in naïve controls (Mann-Whitney, U=26, *p=0.0282). **e)** Experimental design to assess the impact of drug access history on non-contingent CS-evoked dopamine release. Following self-administration acquisition, and an additional week of daily ShA sessions (week 1), subjects were split into short and long-access groups (ShA and LgA, respectively), with ShA continuing with 1-hour access and LgA transitioning to six-hour access, daily for ten additional sessions (weeks 2 and 3). Non-contingent cue probe sessions (red arrows) were interleaved at the end of weeks 1 and 3. Week 1 ShA subjects were the same as those in c. **f)** Mean (± SEM) drug intake during the first hour of self-administration from ShA (open circles) and LgA (closed circles) groups. Blue circles indicate data from ShA sessions, and magenta from LgA sessions. LgA subjects took significantly more drug in the first hour than ShA (Mixed Effects REML-main effect of access: F(1,91) = 8.502, p<0.01; access x time interaction: F(9, 760) = 2.942, §§ p<0.01) **g)** Left: Mean (+SEM) non-contingent CS evoked dopamine response traces from ShA (top) and LgA (bottom) cohorts at week 1 and week 3 timepoints. Right: Mean (+SEM) CS-evoked [Dopamine](nM) from ShA and LgA groups. CS-evoked dopamine responses increased with drug experience (Mixed Effects REML-main effect of week: F(1,15)=10.14, p<0.01; main effect of drug access: F(1,36)=4.297, p<0.05), and this effect is robust in LgA (Sidak post-hoc test **p<0.01). Data from ShA weeks represented in blue, and LgA weeks in magenta. In all cases, n indicates the number of biological replicates (subjects).

We next tested how dopamine, evoked by non-contingent CS presentation, evolves over extended drug use. Following five ShA baseline sessions (week 1), animals were split into two groups, receiving either ShA or six-hour long access (LgA) for ten additional self-administration sessions (weeks 2 and 3), with probe sessions interleaved at the end of weeks 1 and 3 prior to a self-administration session (Figure 1e). Animals in the LgA group took significantly more drug in the first hour than ShA animals (main effect of access: F(1,91) = 8.502, p<0.01; access x session interaction: F(9, 760) = 2.942, p<0.01; Figure 1f) as previously reported^17^. During probe sessions, there was an increase in CS-evoked dopamine between week 1 and week 3, which was significant in the LgA cohort (Sidak post-hoc test: p<0.01; main effect of access: F(1,26)=4.297, p<0.05; Figure1g). These data demonstrate that non-contingent CS-evoked dopamine release increases over the progression of drug use.

### Increased cue-evoked dopamine elevates craving

We next tested whether this increase in cue-evoked dopamine release is modulated by psychological states, specifically cue-induced craving. To manipulate the psychological state without extending drug-intake history, we used a task that models changes in drug seeking during periods of drug abstinence. The frequency that animals perform an action to earn presentation of a previously drug-paired cue increases as a function of time since they last received the drug, a phenomenon termed ‘incubation of craving’^18^. We replicated this phenomenon using a within-animal design^19^ (Figure 2a) where responding for the CS in the absence of cocaine delivery was significantly greater at one month compared to one day following the last drug access (Wilcoxon matched-pairs signed rank test: n=12, W=78, p<0.001; Figure 2b). While this metric evaluates *CS-reinforced* drug seeking (conditioned reinforcement), it is not a direct measure of *CS-elicited* drug seeking. Therefore, we also quantified conditioned approach behavior from video recordings during probe sessions the day before extinction sessions (Figure 2a). We presented the CS five times during probe sessions and found an increase in CS-elicited approach (Wilcoxon matched-pairs signed rank test: n=13, W=86, p=0.001; Figure 2c), demonstrating that cue reactivity also incubates during drug withdrawal. Concomitant to this behavioral incubation in probe sessions, there was a robust increase in CS-evoked dopamine release (Wilcoxon matched-pairs signed rank test: n=7, W=28, p<0.05; Figure 2d).

**Figure 2.**
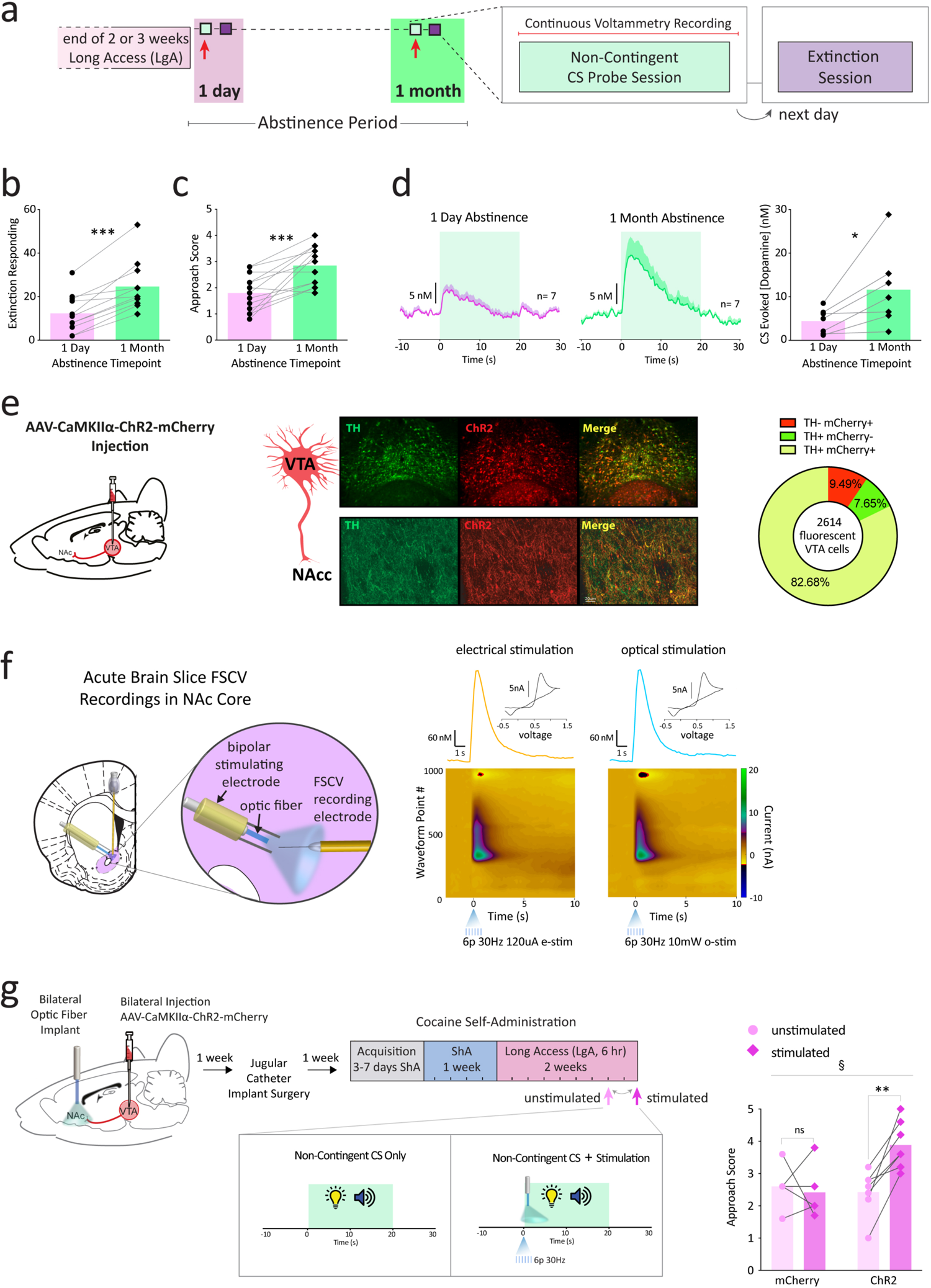
Increased cue-evoked dopamine elevates craving. **a)** Experimental paradigm used to assess CS-evoked drug-seeking behavior and dopamine release during abstinence. Non-contingent CS Probe sessions (red arrow, light green boxes) were conducted 1-day and 1-month following the last LgA session. Five CS presentations were delivered in each probe session, at three minute intervals. Extinction sessions (purple boxes) were carried out one day after each non-contingent probe session. **b)** Extinction responding for the CS alone during abstinence (30 min session). Within-subject repeated measures represented by connected circles (1-day) and diamonds (1-month), with overlaid bars indicating the mean (pink: 1-day, green: 1-month) for b-d, e. Extinction responding significantly increased from 1-day to 1-month of abstinence (Wilcoxon matched-pairs signed rank test: n=12 W= 78, ***p<0.001). **c)** CS-evoked approach behavior during abstinence. Conditioned approach was scored on scale from 0-5, with 0= no response and 5= orient, approach and interaction with active nose poke port. Approach scores significantly increased following 1 month of abstinence (Wilcoxon matched pairs signed rank test: n=13 W=86, ***p=0.001). **d)** Left: Average non-contingent CS evoked dopamine traces from 1-day (pink) and 1-month (green) abstinence probe sessions. Right: Mean (+SEM) CS-evoked [Dopamine] (nM) significantly increased following 1-month of abstinence (Wilcoxon matched-pairs signed rank test: n=7, W=28, *p<0.05). **e)** Left: Rats were injected with AAV1-CAMKIIα-ChR2-mCherry into the ventral tegmental area (VTA). Middle: Fluorescence microscopy imaging of tyrosine hydroxylase (TH) and mCherry expression in VTA cell bodies (top row) and nucleus accumbens (NAcc) axon terminals (bottom row). Right: Selectivity of ChR2 expression in dopamine neurons was estimated by assessing overlap between TH+ and mCherry+ cells. 82% of 2614 cells counted expressed both TH and mCherry. **f)** Left: ChR2 functionality was assessed by measuring optically stimulated dopamine release with FSCV in live brain slices containing the NAcc. Right: Representative pseudo color plots and corresponding dopamine traces and cyclic voltammograms (inset) triggered by optical and electrical stimulation. Pseudo color plots show current changes across the full range of applied voltages (Eapp; y axis) over time (x axis). Optical stimulation (6 pulses, 30Hz, 10mW) of the NAc (right plot) produced a similar dopamine response to an electrical stimulation (left plot) with the same pulse and frequency parameters (6 pulses, 30Hz, 120uA). **g)** Left: To determine the causal influence of non-contingent CS-elicited dopamine release on conditioned approach behavior, the impact of optogenetic stimulation of dopamine release on CS-evoked conditioned approach behavior was assessed in two probe sessions occurring in the second week of LgA in a new cohort of animals. In one session, non-contingent CS presentation onset was paired with brief optogenetic stimulation (6 pulses, 30 Hz, 10 mW; stimulated session, dark pink arrow), and in the other no stimulation was delivered (unstimulated session, light pink arrow). Each probe consisted of five CS presentations, occurring at three-minute intervals. This procedure was carried out in both animals expressing AAV1-CAMKIIα-ChR2-mCherry (ChR2) and AAV1-CAMKIIα-mCherry (mCherry controls). Stimulated and unstimulated probe sessions were counterbalanced across subjects and interleaved by at least two days of self-administration. Right: CS-evoked approach behavior scores from stimulated (dark pink diamonds) and unstimulated (light pink circles) probe sessions, paired data from each subject are connected by lines. Bars indicate the across-subject mean. Conditioned approach was significantly more robust in stimulated sessions in ChR2-expressing animals, but not in controls (virus x stimulation interaction: F(1,10)=8.778, §p<0.05; Sidak post-hoc: significant effect of stimulation in ChR2 group, **p<0.01).

To test causality between these changes in cue-induced phasic dopamine signaling and drug-seeking behavior, we augmented CS-elicited dopamine after one day of abstinence, to mimic the neurochemical changes observed during incubation of craving. Rats received bilateral microinjections into the ventral tegmental area (VTA) of a viral vector containing Channelrhodopsin-2 (ChR2) behind a CaMKIIα promoter (Figure 2e), enabling manipulation of dopamine release with high temporal precision (Figure 2f). Animals were equipped with bilateral optic fibers into the NAcc and underwent cocaine self-administration training. During the final week of LgA, self-administration sessions were interleaved with two counterbalanced non-contingent CS probe sessions, one with CS presentation alone, and one with stimulation (6 pulses, 30 Hz, 10 mW) during CS presentation (Figure 2g). Conditioned approach was significantly more robust in stimulated than unstimulated sessions in ChR2-expressing animals, but not in controls (virus x stimulation interaction: F(1,10)=8.778, p<0.05; Figure 2h), establishing a causal role for dopamine. Together, these data indicate that the progressive elevation of cue-evoked dopamine, across phases of the addiction cycle, increases cue-induced craving.

### Individual drug-taking patterns co-vary with changes in cue reactivity

Thus far, we have treated drug-access groups homogeneously. However, individual differences in behaviors, such as drug consumption rates and craving^20^, are a hallmark of human substance use and can be observed in animal models. Therefore, we tested whether individual differences in the propensity to escalate drug consumption during LgA bore any relation to changes in cue-induced craving. We classified the animals that had undergone LgA as ‘escalators’ or ‘non-escalators’ based upon linear regression of their (first-hour) daily cocaine intake over time, as previously described^15^. 40.9 % (27/66) of animals fulfilled the criterion for escalator classification by progressively increasing their cocaine consumption across sessions, whereas the remaining animals (39/66) exhibited stable drug intake (Figure 3a). When we re-analyzed CS-evoked approach behavior with respect to these groups, we found differential effects over drug-use history (escalation group x week interaction: F(2,18)=4.853, p<0.05), with conditioned approach increasing over time in escalators but not in non-escalators (Figure 3b). Indeed, there was significant correlation between escalation slope and change in CS-evoked approach across individual animals (r^2^=0.47, p<0.05; Figure 3c). Phasic dopamine release to non-contingent CS presentations also changed differentially over time between escalation groups (escalation group x session interaction: F(2,17)=6.546, p<0.01) with dopamine responses increasing in escalators but remaining stable in non-escalators (Figure 3d, e). These changes in CS-evoked dopamine significantly correlated with the concomitant change in CS-elicited approach (r^2^=0.85, p<0.001; Figure 3f), consistent with the causal link between dopamine and conditioned approach that we demonstrated above. These data establish that individuals who escalate their drug intake exhibit a corollary increase in dopamine release evoked by non-contingent cue presentation and consequent behavioral cue reactivity.

**Figure 3.**
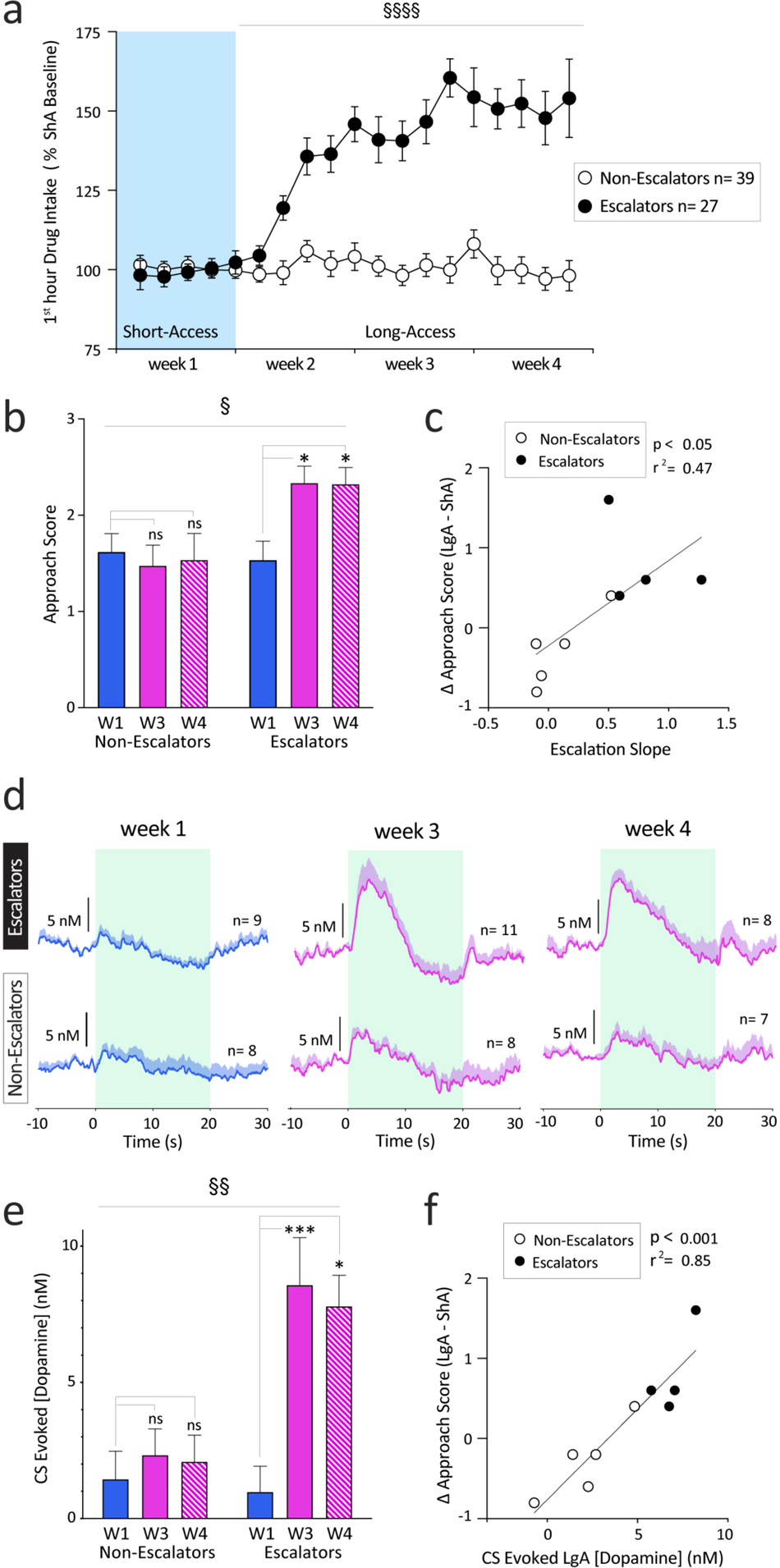
Individual drug-taking patterns co-vary with changes in cue reactivity. Animals in the LgA group from Figure 1, were classified into escalation groups based on their individual drug-taking patterns, as previously established^15^. Animals for which the linear regression of first-hour drug intake over sessions had a significant, non-zero slope, were classified as ‘escalators’, and all others were classified as ‘non-escalators’. **a)** Mean (±SEM) first hour drug intake for non-escalators (open circles) and escalators (closed circled). escalation group x session interaction: F(13,647) = 6.943, §§p<0.0001 **b)** CS-evoked approach behavior measured after week 1 (W1) of ShA and weeks 3 and 4 (W3, W4) of LgA re-analyzed with respect to escalation group. Differential effects over drug-use history were observed between escalators and non-escalators (Mixed Effects REML main effect of week: F(2,18)=3.859, *p<0.05; group x week interaction: F(2,18)=4.853, §p<0.05), with conditioned approach increasing over time in escalators but not in non-escalators (post-hoc Dunnett’s test: for escalators both W1 versus W3 and W4 *p<0.05). **c)** Correlation between escalation slope and change in CS evoked approach behavior (n=9, r^2^=0.47, *p<0.05). Non-escalators and Escalators indicated by open and filled circles, respectively. **d)** Mean (+SEM) non-contingent CS evoked dopamine responses at W1, W3 and W4 re-analyzed with respect to escalation groups: escalators (top) and non-escalators (bottom traces). **e)** Mean (+SEM) CS-evoked [Dopamine](nM) also changed differentially over time between escalation groups (Mixed Effects REML main effect of escalation group: F(1,20)=7.328, *p<0.05; main effect of week: F(2,17)=10.77, ***p=0.001; escalation group x session interaction: F(2,17)=6.546, §§p<0.01), with dopamine responses increasing in escalators but remaining stable in non-escalators (post-hoc Sidak test: escalators W1 versus W3 ***p<0.001 and W1 versus W4 *p<0.05). **f)** Correlation between LgA non-contingent CS evoked [Dopamine](nM) in and change in CS-evoked approach behavior (n=9, r^2^=0.85, ***p<0.001). Non-escalators and escalators indicated by open and filled circles, respectively.

### Diametric dopamine trajectories underlie different aspects of substance use disorders

This increase in dopamine release to non-contingent CS presentation in escalators is surprising given our previous demonstration of phasic dopamine release evoked by the same CS, presented as a result of the animal’s active self-administration (i.e., response-contingent), is attenuated following escalation^15^. To confirm this previous finding, we examined phasic dopamine release during the self-administration sessions that interleaved the CS probe sessions. Similar to our previous work, dopamine release to response-contingent CS presentation significantly declined in escalators but not in non-escalators (escalation group x session interaction: F(2,23)=3.655, p<0.05; Figures 4a, b). Thus, we found that, in animals that escalate cocaine consumption, CS-evoked dopamine release follows opposite trajectories within the same animal, depending on the contingency of the CS; whereas dopamine release evoked by CS presentation is relatively stable in non-escalators for both contingencies. Moreover, we found significant correlation between changes in dopamine and the extent of escalation (r^2^ = 0.30, p<0.05; Figure 4c). To test the causal relationship between these neurochemical and behavioral variables, we once again turned to optogenetic stimulation of dopamine release, time locked to CS presentation. This approach allowed temporally precise stimulation (6 pulses, 30 Hz, 5 ms, 10 mW) of phasic dopamine at the time of response-contingent CS presentation during the first hour of a LgA session (Figure 4d). Stimulation significantly reduced drug consumption in ChR2-expressing escalators, but not in controls, nor in non-escalators (virus x escalation group x stimulation Interaction: F(1, 25)=15.96, p<0.001; Post-hoc Sidak comparison of stimulated versus unstimulated sessions in ChR2 escalators: p<0.0001; Figure 4e). Interestingly, stimulation in ChR2 non-escalators produced a modest *increase* in drug taking (p<0.01; Figure 4e). However, this effect was not selective for animals who received the ChR2 virus as there was not a significant virus x stimulation interaction for the non-escalators (F(1,15)=0.001, p > 0.05; Figure 4e). Therefore, the context of cue presentation dictates the directionality of changes in cue-evoked dopamine amplitude over the course of drug taking in escalators. These changes underlie different aspects of substance use disorders: *Reduced* dopamine to contingent drug-related stimuli produced escalation of drug consumption whereas *increased* dopamine release to non-contingent cue presentation produced elevated craving.

**Figure 4.**
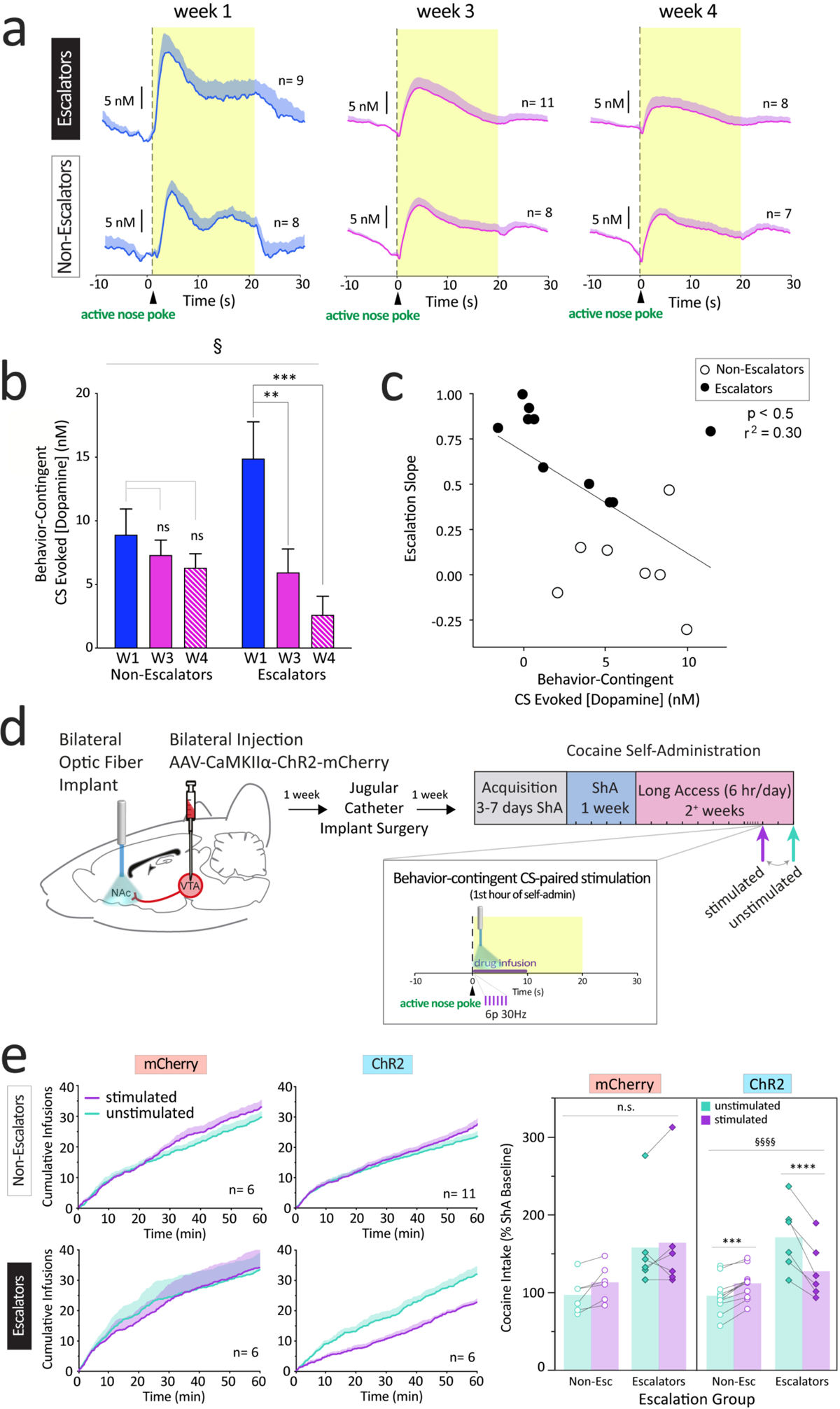
Diametric dopamine trajectories underlie different aspects of substance use disorders. **a)** Mean (+SEM) behavior-contingent CS-evoked dopamine release from escalators (top) and non-escalators (bottom) after Week 1 of ShA (left plot) and Weeks 3 and 4 of LgA (middle, right plots respectively). **b)** Mean (+SEM) response-contingent CS-evoked [Dopamine](nM) significantly declined in escalators, but not non-escalators (Mixed Effects REML main effect of week: F(2,23)=8.59, **p<0.01; escalation group x session interaction: F(2,23)=3.655, §p<0.05; post-hoc Dunnett’s comparison between escalators W1 versus W3 **p<0.01, and W1 versus W4 ***p<0.001). **c)** Correlation between LgA behavior-contingent CS-evoked [Dopamine](nM) and escalation slope. Non-escalators and escalators indicated by open and filled circles, respectively. Significant negative correlation (n=17, r^2^=0.30, *p<0.05). **d)** Experimental design for assessing causal relationship between behavior-contingent CS-evoked dopamine and drug-intake. In the second week of LgA, a new cohort of animals, expressing either ChR2 or mCherry (controls) underwent counterbalanced optogenetic probe tests. In stimulated probes, optical stimulation (6 pulses, 30Hz, 10mW) was paired with each response-contingent CS during the first-hour of self-administration, whereas animals were tethered but no stimulation was delivered in unstimulated probes. **e)** Left: Mean (+SEM) cumulative cocaine intake from stimulated (purple) and unstimulated (cyan) sessions in mCherry (left) and ChR2 (right). Data from non-escalators and escalators are plotted separately (top and bottom, respectively). Right: Cocaine intake during unstimulated (cyan) and stimulated (purple) self-administration sessions from mCherry (left panel) and ChR2 (right panel). Bars indicate mean cocaine intake from unstimulated (cyan) and stimulated (purple) sessions for each escalation group, with individual subject data overlaid (non-escalators: open circles, escalators: filled diamonds, lines connect data from the same subject). Stimulation significantly decreased drug consumption in ChR2-escalators and produced a modest but significant increase in drug consumption in ChR2-non-escalators. Stimulation had no effect on drug intake in mCherry controls. (Repeated Measures 3-Way ANOVA: virus x escalation group x stimulation Interaction: F(1, 25)=15.96, §§§p<0.001. Follow up Repeated Measures 2-Way ANOVA in ChR2 subjects: escalation group x stimulation interaction: F(1,15)=104.8, §§§§p<0.0001; Post-hoc Sidak comparison between stimulated and unstimulated sessions in ChR2 escalators, ****p<0.0001, and ChR2 non-escalators, ***p<0.001. Follow up Repeated Measures 2-Way ANOVA in mCherry subjects: escalation group x stimulation interaction, F(1,10)=0.62, n.s. p=0.45; main effect of stimulation, F(1,10)=3.057, n.s. p=0.1; main effect of escalation group, F(1,10)=3.922, n.s. p=0.076)

## Discussion

Here we demonstrate that dopamine release to non-contingent presentation of drug-related stimuli increases following drug experience and has a causal role in producing drug-seeking behavior. These experience-dependent changes in dopamine differ between subjects. Concomitant increase in neurochemical and behavioral reactivity to drug cues is exhibited by individuals that escalate drug consumption, but not by those that maintain stable drug intake. In individuals that escalate their drug intake, the dopamine signal to response-contingent stimulus presentation during drug taking changes in the opposite direction with experience (i.e., decreases), causing the escalation. Thus, dopamine release to drug-related stimuli is dynamic over the history of drug taking in a subset of individuals, but the directionality of change depends on how the stimulus is encountered. These studies highlight the importance of the stimulus-presentation context in determining and interpreting the neurobiological substrates underlying behavior elicited by drug-associated stimuli.

One conceptual framework on the regulation of drug consumption has linked dopamine to drug satiety, with elevated dopamine levels providing negative feedback on subsequent drug taking^21, 22^. The current data advance this model by demonstrating a causal role for *phasic* dopamine in regulating drug consumption, where discrete bouts of activity consisting of just a few action potentials can have potent inhibition of subsequent drug intake. Moreover, these data identify how decreased dopamine release during drug intake, observed in a subset of animals, produces escalation of drug consumption in those individuals. However, this framework provides no obvious insight into neurochemical changes that take place in response to non-contingent stimulus presentation and their impact on drug craving. In contrast, the central tenet of the incentive sensitization theory is that the release of dopamine following cue presentation attributes incentive value to that stimulus and this signaling is augmented with chronic drug use^7^. The effects we observed related to non-contingent cue presentation phenotypically align well with this theory, providing concrete evidence for several of its critical predictions. Specifically, we demonstrate that cue-evoked dopamine increases over the history of substance use, it does so with the most robust changes in individuals that exhibit the strongest proxies of craving and in contexts that enhance craving, and it is sufficient to elevate craving-like behavior. Nonetheless, we cannot conclude unequivocally that these empirical data are mechanistically explained by the theory as it is still possible that dopamine is operating purely as a teaching signal rather than invigorating the psychomotor activation to the cue presentation *per se*. That is, our data are unable to discern whether dopamine maintains conditioned responding to multiple presentations of the stimulus by directly attributing incentive salience, or by preventing extinction of the association. This distinction is subtle and is seldom tenable from empirical data but is conceptually important for a full understanding of the role that dopamine plays in addiction. Collectively, the current data provide an important perspective into the multidimensional role of dopamine in drug use that is not explained by any single contemporary theory of substance use disorders.

The behavioral model used in this study permits a granular analysis of the neurochemical regulation of substance use, while maintaining relevance to clinical neuroscience. It employs voluntary drug intake where subjects exhibit escalation of consumption with repeated use^17^ which, like with human substance use^20^, is subject to individual differences^15^. Using this approach, we observed systematic changes in drug-cue reactivity, a process that significantly predicts human drug-use and relapse outcomes^5,6^. Importantly, it is well established in humans that drug-paired cues can elicit dopamine release^23^, and produce craving^1–3^. Our data replicate each of these findings, demonstrate how they evolve with substance use, and establish a causal link between them. We also observed a relationship between dopamine release to non-contingent cue presentation and ‘incubation of craving’. This behavioral phenomenon was first observed in humans^24^, formally characterized in rodents^18^ and subsequently revisited in clinical populations where physiological correlates have been successfully measured^25^. A key clinical implication of the current results reinforces the importance of tailoring potential therapeutics for substance use disorders to the phase of drug use. Treatments that have direct or downstream effects on dopamine release should be better suited to combating either active drug use or relapse depending on whether they increase or decrease dopamine transmission, respectively. Accordingly, clinical studies with levodopa (which increases dopamine), show that its efficacy depends on the baseline status of the patient, with promising effects only on those with active drug use at the start of treatment^26^. Therefore, the current findings dovetail with existing literature in human subjects and, through the benefits of a model system, are able to provide several unique insights into the neurobiological regulation of substance use.

Overall, we have demonstrated that phasic dopamine release evoked by drug-related stimuli changes dynamically over the course of cocaine use in a subset of subjects. These individuals exhibit consequential behaviors that model core symptoms of substance use disorders. Remarkably, the changes in dopamine signaling are diametrically opposed between substance-use contexts. Dopamine release to drug cues encountered non-contingently increases to produce elevated cue-evoked craving. Whereas dopamine release to stimuli presented as a result of drug-seeking behavior decreases, conferring increased drug consumption.

## Methods

### Subjects

A total of 217 adult male Wistar rats (Charles River, Raleigh NC) weighing 300-350g were used in these studies. Rats were housed individually and kept on a 12-h light/12-h dark cycle (lights on at 0700) with controlled temperature and humidity, and food and water available *ad libitum*. All animal use was approved by the University of Washington Institutional Animal Care and Use Committee, and surgical procedures were performed under aseptic conditions.

Twenty-Seven rats completed behavioral training in the short-access (ShA cohort) and 66 in the long-access (LgA) cohort. Successful voltammetry recordings were obtained from at least one experimental timepoint in 15 ShA animals, and 30 LgA animals. Thirteen subjects that completed LgA subsequently completed incubation of craving behavioral studies during abstinence, and voltammetry recordings were obtained from seven of these animals. Thirty-four rats completed the optogenetics studies. Approximately 20% of subject attrition was due to head cap loss or catheter failure during the post-surgery recovery period prior to beginning experimentation. The remainder of subjects dropped out of the study after experimentation began due to head cap loss, electrode failure, catheter failure or rejection, lack of viral expression, or in rare instances, failure to acquire self-administration. In all cases, the number of subjects reported, n, equals the number of biological replicates (subjects) included in the dataset.

### Data and Code Availability

All data and code used in the manuscript will be made available in a public repository from acceptance of the manuscript.

### Surgery

For dopamine recording studies, chronically implantable carbon fiber microelectrodes, constructed as previously described^27, 28^, were lowered unilaterally or bilaterally into the nucleus accumbens core (NAcc)^29^, (AP: +1.3mm, ML: ±1.3mm, DV: −7.2mm) using a stereotaxic frame, and secured with dental acrylic. After two weeks of post-operative recovery, rats underwent a second surgical procedure and were outfitted with indwelling intravenous jugular catheters, then allowed to recover for at least one week before initiating cocaine self-administration training. During this recovery period, and prior to days off from self-administration, catheters were backfilled with a viscous 60% polyvinylpyrrolidone-40 (Sigma Aldrich, USA) solution, containing 20 mg/mL of gentamicin (40 mg/mL; Fresenius Kabi, USA), and 1,000 IU/mL of heparin (20,000 IU/mL; Fresenius Kabi, USA) to prevent formation of blood clots. Otherwise, catheters were flushed daily with saline or 80 IU/mL of heparinized (1,000 IU/mL; Fresenius Kabi, USA) saline as needed, to maintain catheter patency throughout experimentation.

For optogenetic studies, rats first underwent catheter implantation surgery (as described above), and following one week of recovery underwent intracranial surgery in which they were bilaterally injected with AAV1-CAMKIIα-ChR2-mCherry or AAV1-CAMKIIα-mCherry (viral vectors were made and provided by Dr. Larry Zweifel, Univ. of Washington) into the VTA^29^ (AP: −6.35mm, ML: ±0.5mm, DV: −8.5mm) and optical fiber stubs (1.25mm stub diameter, 200μm fiber diameter, Plexon Inc.) were implanted bilaterally into the NAcc^29^ (AP: +1.3mm, ML: ±1.3mm, DV: −7.0mm). After an additional week of recovery, rats began cocaine self-administration training. A minimum of three weeks elapsed between viral vector injections and optogenetic manipulation to allow for ample ChR2 expression.

### Cocaine Self-Administration

Rats were trained to self-administer cocaine during daily one-hour (short-access, ShA) sessions in an operant chamber outfitted with a liquid swivel and containing two nose-poke ports. During self-administration sessions, the illumination of a house light paired with white noise signaled the availability of drug. A single nose poke into the active port elicited a 0.5 mg/kg cocaine infusion (fixed-ratio one schedule), accompanied by presentation of a 20 second audiovisual conditioned stimulus (CS, nose-poke light + tone), during which any additional nose-poke was without consequence (time out). Nose pokes into the inactive port at any time were without consequence. After meeting the acquisition criterion of performing three sequential sessions where 10 or more infusions were earned, animals received five additional daily ShA sessions to establish baseline intake. Animals were then divided into two drug access groups, each receiving the same number of sessions but sessions differed with respect to the number of hours the animals had access to self-administer cocaine. The short-access (ShA) cohort received daily one hour access for 10 additional days, while the long access (LgA) group received six-hour access for 10 days. A subset of LgA animals received an additional five sessions (mirroring the duration of LgA used in our previous studies^15^). To assess individual differences in drug intake patterns observed in LgA animals, we used our previously validated method^15^ for separating escalators and non-escalators. We conducted a linear regression predicting each animal’s first-hour drug intake across sessions (post-acquisition), and rats having a significant, positive non-zero slope were considered escalators.

### Assessment of Drug-Seeking Behavior

In drug-free, non-contingent CS probe sessions, conditioned approach behavior elicited by five non-contingent CS presentations was assessed and CS-evoked dopamine release was measured. Conditioned approach was scored on a 0-5 scale with the following criteria: 0 = no response; 1= startled action to cue onset; 2 = animal’s head directs towards nose-poke port; 3 = animal orients body towards nose-poke port; 4 = animal orients body and approaches nose-poke port; 5= animal orients, approaches, and interacts with the nose-poke port. In cases where the animal’s head was outside of the camera view, data from the trial was excluded from statistical analysis. Non-contingent CS probe sessions occurred once per week (excluding week two) immediately prior to drug self-administration sessions, or on their own during a period of abstinence following drug use at one day and one month abstinence timepoints.

CS-reinforced drug-seeking was measured in separate, thirty-minute extinction sessions occurring one day after non-contingent probe sessions during abstinence. These sessions were identical to self-administration sessions, except that nose-pokes into the active port elicited the CS alone, without the delivery of drug (infusion pumps were off and infusion lines were backfilled with saline).

### Fast-Scan Cyclic Voltammetry

Behaviorally relevant stimuli elicit rapid changes in dopaminergic neuron firing, resulting in transient changes in dopamine release over the course of seconds, and requiring the use of a detection method with high temporal resolution. In the studies phasic dopamine release events were measured at carbon fiber microelectrodes in the ventral striatum using fast-scan cyclic voltammetry (FSCV)^27, 28^. Briefly, chronically implanted carbon-fiber microelectrodes were connected to a head-mounted voltametric amplifier, interfaced with a PC-driven data-acquisition and analysis system (National Instruments, TX, USA) through a commutator (Med Associates, VT, USA) that was mounted above the test chamber. A potential was applied to the electrode as a triangular waveform such that it was linearly ramped from the initial holding potential (−0.4 V vs Ag/AgCl) to a maximum voltage (1.3 V vs Ag/AgCl, anodic sweep), then returned to the holding potential (cathodic sweep). Each voltage scan lasted 8.5 ms, yielding a scan rate of 400 V/s. The holding potential was maintained between voltage scans. Scans were applied every 100 ms (10 Hz sampling). When dopamine was present at the surface of the electrode, it was oxidized during the anodic sweep to form dopamine-*o*-quinone (peak reaction detected at approximately +0.7 V), which was reduced back to dopamine in the cathodic sweep (peak reaction detected at approximately −0.3 V). The ensuing flux of electrons was measured as current and was directly proportional to the number of dopamine molecules that underwent electrolysis. Voltametric data was band-pass filtered at 0.025–2,000 Hz. The background-subtracted, time-resolved current obtained from each scan provided a chemical signature characteristic of the analyte, allowing resolution of dopamine from other substances^30^. Dopamine was isolated from the voltametric signal by chemometric analysis using a standard training set^27^ based on electrically stimulated dopamine release detected by chronically implanted electrodes, and dopamine concentrations estimated on the basis of the average post-implantation sensitivity of electrodes. Before analysis of average concentration, all data were smoothed with a five-point within-trial running average. The mean concentration of dopamine was obtained by averaging over the seven seconds (approximate duration of the observed phasic signal) following the non-contingent presentation of the CS or operant response to obtain drug. Data collected on recording days was included when there was a detectable dopamine response at any point within the session. Data was excluded from individual sessions where electrical noise exceeded 0.2 nA, animals became disconnected from the drug delivery tubing, or when tethering for recordings altered the animal’s regular behavior patterns. In the rare cases where more than one electrode was functioning in an animal during a given session, the average of the signals obtained from both electrodes was used for analysis. In cases where we only obtained voltammetry data from a single experimental time point (session) from an animal, this data was excluded to minimize any electrode sensitivity bias.

### Optogenetics

To determine the causal influence of non-contingent CS-elicited dopamine release on conditioned approach behavior, we performed optogenetic manipulation of dopamine release during non-contingent probe sessions in the second week of LgA. In one session, non-contingent CS presentation onset was paired with brief optogenetic stimulation with a 465nM LED (6 pulses, 30 Hz, 5 ms pulse width, 10 mW; Plexbright Compact LED, Plexon Inc.; stimulated session), and in the other no stimulation was delivered (unstimulated session). This procedure was carried out in both subjects expressing AAV1-CAMKIIα-ChR2-mCherry (ChR2) and AAV1-CAMKIIα-mCherry (mCherry controls). Stimulated and unstimulated probe sessions were counterbalanced across subjects and interleaved by at least two days of self-administration. Non-contingent CS-elicited approach behavior was scored as described above and compared between stimulated and unstimulated sessions.

To determine how behavior-contingent CS-elicited dopamine release influenced drug-intake, we paired optogenetic stimulation of dopamine release with the behavior-contingent CS presentations occurring during the first hour of drug self-administration. This manipulation was limited to the first hour to minimize carryover effects on drug-taking behavior. As in the previous study, both ChR2 and mCherry control subjects underwent counterbalanced test sessions (interleaved by at least two days of self-administration). In one session, stimulation was paired with response-contingent CS (6 pulses, 30 Hz, 5 ms pulse width, 10 mW; stimulated session), and another without stimulation (unstimulated). The number of drug-infusions earned in the first hour was recorded and comparison within subjects made between stimulated and unstimulated sessions.

### Statistics

Simple comparisons between two conditions were performed using a Mann-Whitney U test or a Wilcoxon matched-pairs signed rank test for between- and within-subjects designs, respectively. Analysis of correlations between variables were conducted using linear regression. Separate slopes and intercepts for each group in the model were examined, and only a single slope and intercept parameter model was fit to all data if no significant group differences were found. In the case of analyses between multiple groups with repeated-measures, ANOVAs were performed using a linear mixed effect modeling approach to account for missing data at random. The model was estimated using restricted Maximum Likelihood (REML), a compound symmetry covariance matrix, and included subjects as a random intercept. When relevant, a maximal random effects structure was used for repeated-measures31. Family-wise error rate of post-hoc simple effects was controlled using either a Holm-Sidak correction, or Dunnett’s test for multiple comparisons with a control condition. All statistical tests were conducted using GraphPad Prism (version 9 for Mac OS, GraphPad Software, www.graphpad.com).

### Histology

Upon completion of experimentation, placements of electrodes (Supplemental Figure 1), or optic fibers (Supplemental Figure 2) and viral expression patterns were assessed. Animals were anesthetized with an intraperitoneal injection of ketamine (100 mg/kg) and xylazine (20 mg/kg) and transcardially perfused with saline then 4% paraformaldehyde. Brains were removed and postfixed in paraformaldehyde for at least 24 h, then serially transferred into 15% and 30% sucrose solutions before being frozen, and coronal sections were taken using a cryostat (Leica CM1850, Leica Inc.) held at −25C. In subjects with electrodes, recording sites were marked by electrolytic lesion prior to perfusion while under anesthesia. Brains from voltammetry studies were sectioned at 50μm thickness and stained with cresyl violet to aid visualization of anatomical structures and electrolytic lesions. Brains from optogenetic studies were sectioned at 40 μm and viral expression patterns and selectivity for dopamine neurons was assessed by immunohistochemistry. In some cases, placements of electrodes or optic fibers could not be obtained because lesions were not clearly visible, or animals lost head caps before perfusion.

### Immunohistochemistry

Brain slices were treated with a blocking solution (phosphate buffered saline containing 3% normal donkey serum and 0.3% Triton) for 1 hour and incubated overnight at 4°C with primary antibodies to label cells containing tyrosine hydroxylase (anti-tyrosine hydroxylase monoclonal, 1:1000 Millipore, MAB318) and mCherry (anti-dsRed polyclonal, 1:1000; Clontech, 632496). Sections were then washed in phosphate buffered saline (PBS) three times for ten minutes each, and incubated for two hours at room temperature with fluorescent conjugated secondary antibodies (1:250 Alexa Fluor® 488 AffiniPure Donkey Anti-Mouse IgG (H+L), 1:250 Cy™3 AffiniPure Donkey Anti-Rabbit IgG (H+L); Jackson Immunoresearch). Sections were washed in PBS three times for ten minutes each, slide mounted and cover slipped with DAPI Fluoromount-G (SouthernBiotech). Fluorescent images of VTA cell bodies were taken using Keyence BZ-X710 (Keyence, Ltd., UK) under 10x magnification, and NAcc terminals under 60x magnification to assess virus and tyrosine hydroxylase expression, and optic fiber placement. Fluorescent cell counting was performed using ImageJ on a total of six coronal slices containing the VTA, two slices from n=3 separate subjects.

## Acknowledgements

We thank Daniel Hadidi, Suhjung Janet Lee, and Hope C. Willis for technical support; and Andy Hart, Yavin Shaham, and Benjamin Saunders for feedback. This work was supported by awards from the US National Institutes of Health grants T32-DA007278 (L.M.B. & R.D.F.), F31-DA048562 (R.D.F), R01-DA039687 (P.E.M.P), R37-DA051686 (P.E.M.P), R01-DA044315 (L.S.Z).

## Author Contributions

L.M.B, R.D.F., I.W. and P.E.M.P. designed research. L.M.B., S.B.E, I.W., S.G.S., L.S.Z., and P.E.M.P. developed the methods. L.M.B, R.D.F., and N.L.M performed *in vivo* recording and optogenetics experiments. L.M.B, and J.S.S. performed slice voltammetry experiments. L.S.Z. designed and made viral constructs. L.M.B, N.L.M., R.D.F., and M.E.S. performed histological analysis. L.M.B, R.D.F., and M.C.P. carried out data analysis. L.M.B. prepared the figures. L.M.B, R.D.F., and P.E.M.P. wrote the initial draft. L.M.B, R.D.F., M.C.P., S.G.S., and P.E.M.P. reviewed and edited the manuscript.

