## Supplemental figures for "Cocaine Seeking And Taking Are Oppositely Regulated By Dopamine"

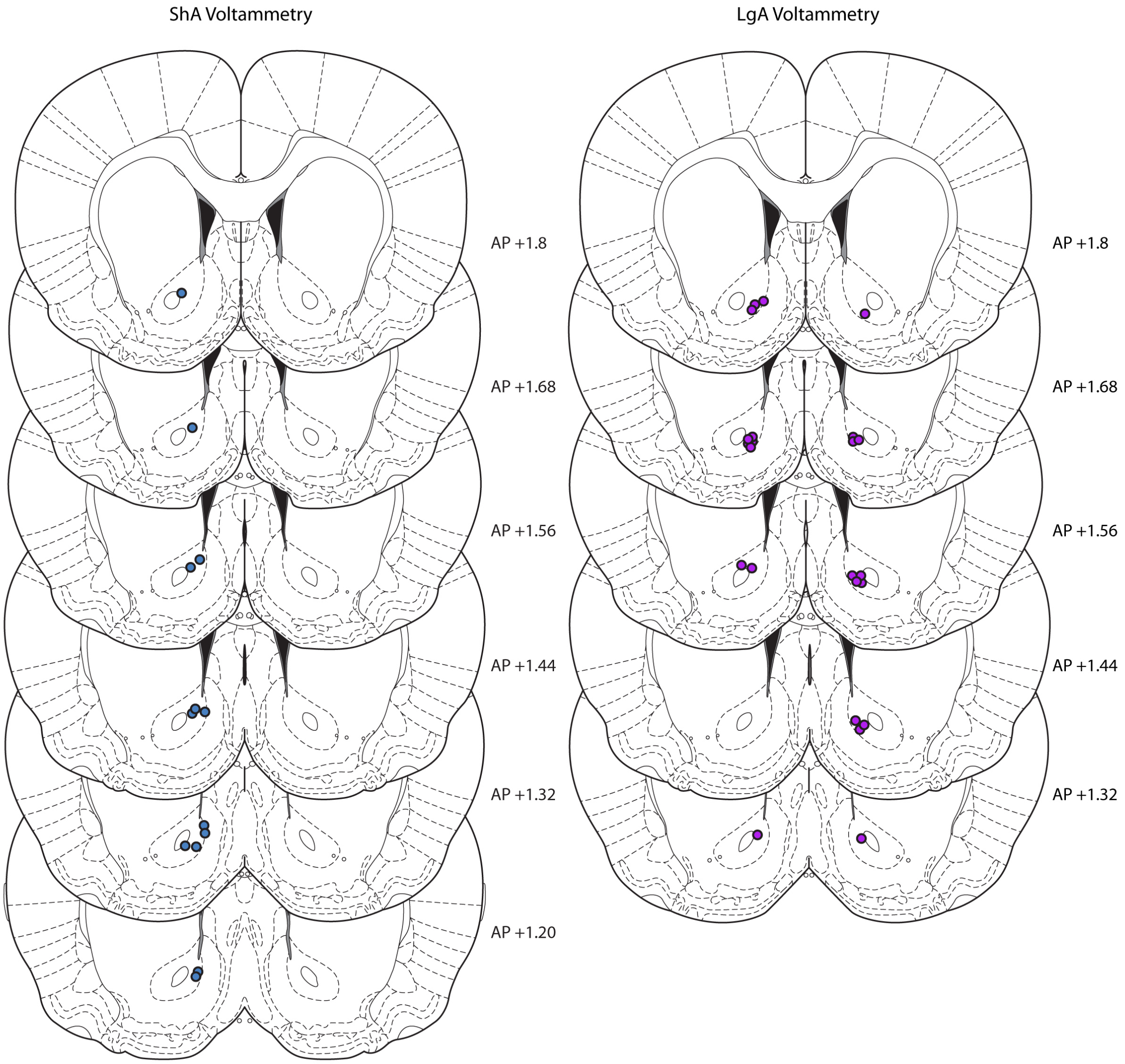


***Supplemental Figure 1. Voltammetry recording sites***

Histological verification of carbon fiber microelectrode placements from voltammetry recording study subjects. a) Electrode placements from ShA cohort subjects indicated by blue circles. b) Electrode placements from ShA cohort subjects indicated by magenta circles. Coronal sections are labeled with anterior-posterior coordinates[29]. Electrode placements for some subjects could not be determined because the lesion was undetectable, animals lost head caps before the end of the study, or brain tissue was damaged during processing.

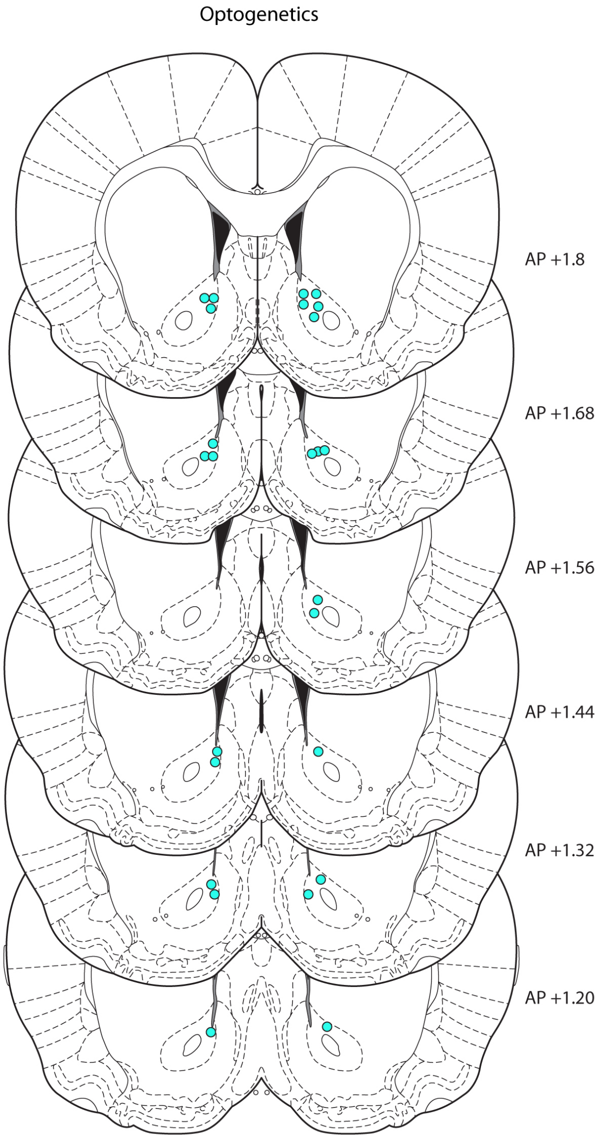


***Supplemental Figure 2. Optical fiber placements***

Histological verification of optic fiber placements are indicated by cyan circles. Coronal sections are labeled with anterior-posterior coordinates[29]. In some cases placements could not be obtained because the fiber track was undetectable, or brain tissue was damaged during processing.
